# An interaction between the transmembrane domains of *Streptococcus pyogenes* sortase A and its endogenous substrate M protein revealed by molecular dynamics simulations

**DOI:** 10.1101/2025.08.19.671115

**Authors:** Nathan G. Avery, Elise F. Tahti, Paul Clinton Spiegel, John M. Antos, James McCarty, Jeanine F. Amacher

## Abstract

Sortase enzymes are cysteine transpeptidases at the cell surface of gram-positive bacteria. Localized to distinct foci on the cell membrane, class A sortases (SrtAs) recognize a cell wall sorting signal (CWSS), and following cleavage at this specific binding motif, target proteins are ligated to precursors of the growing peptidoglycan layer. This activity of SrtA enzymes is utilized extensively in sortase-mediated ligation (SML) strategies, for a variety of protein engineering applications. Typically, engineered variants of SrtA are used for SML experiments considering the relatively low catalytic efficiency of this enzyme. Understandably, most biochemical studies are conducted with the isolated catalytic domain of SrtA enzymes from various bacteria, and the stereochemistry of the endogenous interaction between SrtA and substrate is not well understood. Here, we used AlphaFold2 to create a model of the full-length SrtA enzyme from *Streptococcus pyogenes* (spySrtA) with or without either a peptide substrate or a portion of M protein, a cellular target. We ran triplicate 500 ns molecular dynamics simulations for each model embedded in a lipid bilayer, which revealed several stereochemical features of this system. Contact map analyses revealed specific interactions between catalytic domain positions of spySrtA and the lipid bilayer, as well as between the enzyme and M protein residues outside the canonical LPXTG pentapeptide CWSS. We also characterized a potential transmembrane domain interaction between spySrtA and M protein that we predict orients and stabilizes substrate binding. Taken together, these interactions likely increase the catalytic efficiency of the enzyme for its substrates *in vivo*, and may provide important stereochemical insights for SML uses.

## Introduction

The surface of gram-positive pathogenic bacteria are extensively decorated in proteins.^1^ These include toxins, environmental sensors, components of pili, and proteins with myriad other functions critical to the survival and pathogenicity of these organisms.^1–4^ One mechanism by which covalent attachment of cell surface proteins is achieved includes ligation mediated by sortase enzymes. The first sortase identified was the class A sortase (SrtA) from *Staphylococcus aureus* over 25 years ago.^5,6^ These enzymes are localized to discrete foci on the cell membrane, often the cleavage furrow of dividing bacteria, ligating substrates to precursors of the growing peptidoglycan layer.^7,8^

The catalytic mechanism of SrtA enzymes is well understood. SrtA recognizes a specific cell wall sorting signal (CWSS), defined by the sequence LPXTG, where X=any amino acid and positions are referred to as P4=Leu, P3=Pro, P2=X, P1=Thr, and P1’=Gly.^2,9,10^ Initial cleavage between the P1/P1’ position occurs following nucleophilic attack of the P1 Thr carbonyl carbon by the thiol side chain of a catalytic cysteine residue, forming an acyl enzyme intermediate.^9,10^ A second nucleophile, often the α-amine of an N-terminal amino acid, resolves this intermediate, and the ligation product is formed.^9,10^ In addition to the Cys (C184), the catalytic residues were traditionally thought to include His (H120) and Arg (R197); however, recent work from ourselves and others suggested that while critical, the Arg may not play a catalytic role in electrostatic stabilization of the oxyanion tetrahedral intermediate. We and others determined that this stabilization is instead facilitated by the hydroxyl group of a highly conserved Thr immediately N-terminal to the catalytic Cys, as well as the backbone amide of the amino acid following the catalytic His.^10–12^

The ability to recognize the CWSS followed by specific proteolytic cleavage and ligation of two sequences makes SrtA enzymes versatile tools in sortase-mediated ligation (SML) protein engineering applications.^10,13,14^ *Staphylococcus aureus* SrtA, the first sortase discovered, continues to see regular use for SML experiments. However, because of the strict specificity of this enzyme for the LPXTG target motif, as well as its relatively low catalytic efficiency (a characteristic of all sortases studied to date), engineered variants are most often utilized.^15,16^ Specifically, directed evolution studies identified a pentamutant that increased the catalytic efficiency >100-fold, from 200 M^-1^ s^-1^ (*k*_cat_=1.5 s^-1^, *K*_m_=7.6 mM, for an LPETG peptide) to 23,000 M^-1^ s^-1^ (*k*_cat_=5.4 s^-1^, *K*_m_=0.23 mM).^15^ Despite these advances on the use of sortases *in vitro*, as well as nice early work in the field identifying and investigating the sortase reaction *in vivo*, the role of the sortase transmembrane domain in the catalytic mechanism is not well understood from a biochemical perspective.^6,17–20^

In this work, the increased capabilities of structural modeling, e.g., due to AlphaFold and RoseTTAFold, have allowed us to investigate the behavior of a full-length SrtA enzyme in unprecedented atomic detail for the first time.^21–23^ Specifically, we used AlphaFold2 to model full-length *Streptococcus pyogenes* SrtA (spySrtA), followed by molecular dynamics simulations of this structure in a lipid bilayer mimicking the composition of that in gram-positive bacteria.^24,25^ We chose to investigate spySrtA and not *Staphylococcus aureus* SrtA for several reasons. *Staphylococcus* SrtA enzymes are the only identified that require allosteric activation by calcium,^2,9,10^ and we reasoned that using a non-*Staphylococcus* enzyme may both simplify the overall system and also be more applicable to the superfamily at large. In addition, we recently solved structures of a catalytically inactive variant of spySrtA with peptide substrates (sequences LPATA and LPATS, PDB IDs 7S4O and 7S51) as well as a product mimic (LPAT-LII, PDB IDs 7T8Y and 7T8Z).^26^ To our knowledge, ours are the only experimental sortase structures that contain a non-covalently bound ligand and which show the substrate in the active site conformation that is consistent with known biochemical data.^10^ However, because our structures were deposited in the Protein Data Bank in 2022, and the AlphaFold training database only includes structures deposited in 2021 and earlier,^22^ these act as a structural control for the computationally-generated output models. Finally, although *S. aureus* SrtA and its derivatives are the most widely used enzymes for SML, spySrtA is also utilized, particularly in dual-labeling strategies, due to a larger degree of substrate promiscuity (e.g., at the P1’ position).^10,14,27–29^

Triplicate molecular dynamics simulations were performed, and a number of analyses were employed to better understand the stereochemistry of full-length spySrtA in the membrane, as well as its interaction with an endogenous substrate, M protein. M protein is a well-studied virulence factor in *S. pyogenes* that binds to host proteins and interferes with the host immune response, including by inhibiting phagocytosis.^30^ Specifically, we were curious in characterizing how the CWSS properly binds the active site of spySrtA considering this sequence (LPSTG in M protein) ends very close to the predicted transmembrane domain. We predicted that the lipid bilayer would facilitate positioning of the spySrtA catalytic domain and that the transmembrane domains of both proteins may directly interact. Our MD simulation results suggested that the transmembrane domains of spySrtA and M protein likely do interact, and that specific contacts with lipids may orient spySrtA for catalysis. Furthermore, we identified specific interactions between spySrtA and residues beyond the CWSS, which may play a role in target specificity. Taken together, we predict that the membrane plays a major role in sortase biology *in vivo* and may provide additional insights for further development of these important enzymes for SML engineering applications.

## Materials and Methods

### Structural modeling using AlphaFold2 and software for structural analysis

The sortase A and substrate sequences for structural modeling obtained from Uniprot include: spySrtA (Uniprot ID: Q99ZN4_STRP1) and M protein (Uniport ID: M6A_STRP6). The full-length spySrtA protein sequence was used, and the C-terminal portion of M protein, residues 376-415. Structural models were determined with AlphaFold2 on the European Galaxy Server with default settings.^22,31,32^ AlphaFold2 input sequences can be found in the Supporting Information. Output structures are ranked using the predicted long-distance difference test (pLDDT) and the ranked_0 structure was used for molecular dynamics simulations. Electrostatic potential was calculated using the ABPS plugin and visualized in PyMOL (Schrodinger Software).^33^

### Molecular dynamics simulations

*Streptococcus pyogenes* sortase A (spySrtA) bilayer positioning was calculated using the PPM 2.0 web server using data from the OPM database.^34^ SpySrtA in isolation, with substrate, or peptide was embedded in an 80% 1,2-dioleoyl-sn-glycero-3-phosphoglycerol (DOPG), 20% tetraoleyl-cardiolipin (TOCL2) bilayer using CHARMM-GUI,^35–38^ with CHARMM36m all atom force field.^39,40^ Each system was solvated using TIP3P water and sodium and chloride ions to neutralize the system with an ionic strength of 0.15 M and equilibrated using GROMACS 2022.4 (**Table S1**).^41^ A steepest decent energy minimization was performed until the maximum force on any atom is less than 1000 kJ/mol/nm. The temperature was first equilibrated at 300K with restraints on all heavy protein and lipid atoms using a Berendsen thermostat for 250 ps.^42^ The pressure of the system was equilibrated in the NPT ensemble at 1 bar with decreasing restraints on all heavy protein and lipid atoms using a Berendsen semi-isotropic barostat for 1.75 ns total.^42^ The temperature and pressure of the system was further equilibrated in the NPT ensemble with a Parrinello-Rahman semi-isotropic barostat^43^ and Nose-Hoover thermostat^44,45^ at 300K without restraints for 10 ns. Hydrogen atoms were restrained with the LINCS algorithm.^46^ The equilibrated structures were used to run a single 500 ns simulation at 300K and 1 bar using GROMACS 2022.4.^41^ Simulations were performed in triplicate, with separate equilibrations run for each. Atomic coordinates were saved every 100 ps. The size of each system is reported in **Table S1**.

### Analysis of MD simulations

Contacts between atom pairs (spySrtA, DOPG, TOCL2, and/or substrate) were calculated with PLUMED 2.4.^47,48^ A contact was considered formed if the distance between atoms was less than 4 Å. The contacts included nitrogen, oxygen, and carbon atoms for each residue in spySrtA near the membrane surface and nitrogen, oxygen, and carbon atoms for each lipid type in the membrane (DOPG and TOCL2). Contacts were measured between hydrophilic (nitrogen and oxygen) and hydrophobic (carbon) atoms in spySrtA and each lipid type, and/or between nitrogen, oxygen, and carbon atoms for each residue in spySrtA and substrate. Contacts were monitored over the 500 ns simulations.

Root mean squared fluctuation (RMSF) per residue of backbone atoms, root mean squared deviation (RMSD) of backbone atoms, and solvent accessible surface area (SASA) per residue for side chain and backbone atoms were calculated using GROMACS analysis tools.^49^ Potential energy between spySrtA and substrate or peptide was calculated using GROMACS.

### Bioinformatics analysis of natural spySrtA substrates and homologous Sortase A enzymes

Predicted natural substrates in the *Staphylococcus pyogenes* genome were identified from a Hidden Markov Model (HMM).^50,51^ Five residues before and twenty-three residues after the LPXTG recognition motif sequences were aligned. A logo map of amino acid residue prevalence in each position of the substrate was created using the online WebLogo tool.^52^ A NCBI blast search was performed on the wild-type (WT) spySrtA sequence filtering for the genus *Streptococcus*. Twenty-eight sequences from the *Streptococcus* genus were aligned with spySrtA using Clustal Omega.^53^ Conserved residues were identified using EndScript Server 3.0^54^ and ConSurf.^55,56^ SpySrtA conserved residues with 300 sortase A sequences were identified with ConSurf. Conserved M protein substrate residues were also identified with ConSurf using 300 sequences as an input in the online server. Conservation scores were mapped on to the respective protein structures.

## Results

### The spySrtA catalytic domain interacts with the gram-positive bacterial membrane via electrostatic and hydrophobic interactions

To get a better understanding of the structure of the full-length spySrtA enzyme, we utilized AlphaFold2 modeling, as described in the Materials and Methods (**Figure 1A**). The structure largely agreed with our predictions of the full-length protein; however, we were intrigued by a number of intramolecular contacts in residues of the spySrtA extracellular domain, which are not typically included in the catalytic domain constructs that have been utilized for *in vitro* work (e.g., amino acids 81-249) (**Figure 1B**). Specifically, we observed multiple hydrophobic (Y49-V94-F110, V51-V94, I59-I95) and polar (N47-S243, Q50-N106, S55-E17) interactions (**Figure 1B**). Catalytic domain constructs of sortase A enzymes were historically determined via similarity in multiple sequence alignments, which is how the construct boundaries of spySrtA were defined.^57,58^ However, there is evidence that residues N-terminal to the catalytic domain may play a regulatory role in SrtA enzymes such as *Bacillus anthracis* SrtA.^59^ Moving forward, we will refer to the *extracellular* domain of spySrtA as amino acids 3 4-249 (^34^NKPIR… NQVST^249^) and the *catalytic* domain as amino acids 81-249 (^81^SVLQA… NQVST^249^).

**Figure 1.**
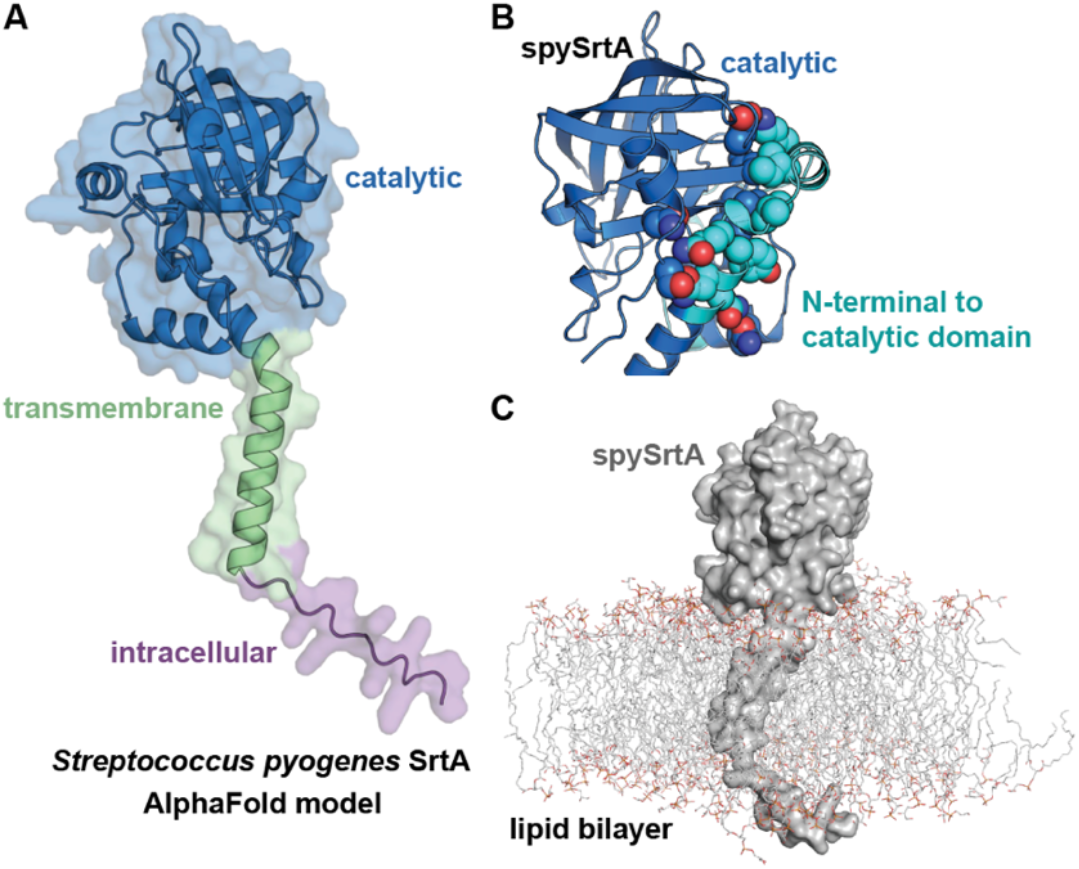
AlphaFold models of full-length *Streptococcus pyogenes* SrtA (spySrtA) with and without a lipid bilayer. (**A**) An AlphaFold2-generated model of full-length spySrtA (residues 1-249) is shown in cartoon representation and including a transparent surface. Three regions of the proteins are colored and labeled. (**B**) Amino acids which may facilitate intraprotein interactions between the catalytic domain (as commonly used in biochemical studies, residues 81-249) and residues N-terminal to this region are highlighted with the side chains shown as spheres and colored by heteroatom (O=red, N=blue, C=marine (for catalytic domain) and C=cyan (for N-terminal to catalytic domain)). SpySrtA is shown in cartoon. (**C**) A full-length model of spySrtA (gray, cartoon representation) in a lipid bilayer (lines, colored by heteroatom with C=gray).

We next modeled spySrtA in its membrane environment, utilizing a lipid composition of 80% 1,2-dioleoyl-sn-glycero-3-phosphoglycerol (DOPG) and 20% tetraoleoyl-cardiolipin (TOCL2), based on previous studies of gram-positive bacterial membranes (**Figures 1C, S1**).^24^ Insertion of spySrtA into the lipid bilayer is described in the Materials and Methods. Following generation of our model of full-length spySrtA enzyme in a lipid bilayer, we ran triplicate molecular dynamics simulations for 500 ns to assess sortase-membrane interactions, as described in the Materials and Methods. Overall, the catalytic domain of spySrtA remained stable during the course of each simulation (**Figure S2**). Contact analyses of specific spySrtA atoms with the lipid bilayer revealed a number of interactions in the catalytic (extracellular) domain that frequently occurred duri ng the simulations (**Figure 2**). As expected, transmembrane residues remained embedded in the lipid bilayer during the entire simulation (**Figures 2A-B**). In addition, residues close to the transmembrane domain were also frequently associated with the membrane exterior. Interestingly, there were also a number of residues not immediately adjacent to the transmembrane domain that were frequently (defined as >50% of the simulation) in contact with lipid carbon or oxygen atoms. These included residues N-terminal to the catalytic domain (S78-E80) or immediately within it (L83, Q86, M87), as well as I147 and T148 near the C-terminus. Some of these residues appeared to preferentially bind to either DOPG or TOCL2. Residues near the peptide binding groove (R38, S78-E80, and L83) bound to the phosphate and/or fatty acid tail of DOPG whereas residues opposite to the peptide binding groove (T40-L41, R44, N47-K48, and Q86-M87) bound to polar and/or hydrophobic groups of TOLC2 for >80% of the simulation (**Figures 2A-B**). Residue-specific preferences for lipid moieties could support the orientations spySrtA adopts in the membrane. Taken together, these data revealed interactions between spySrtA, including the catalytic domain, and the membrane.

**Figure 2.**
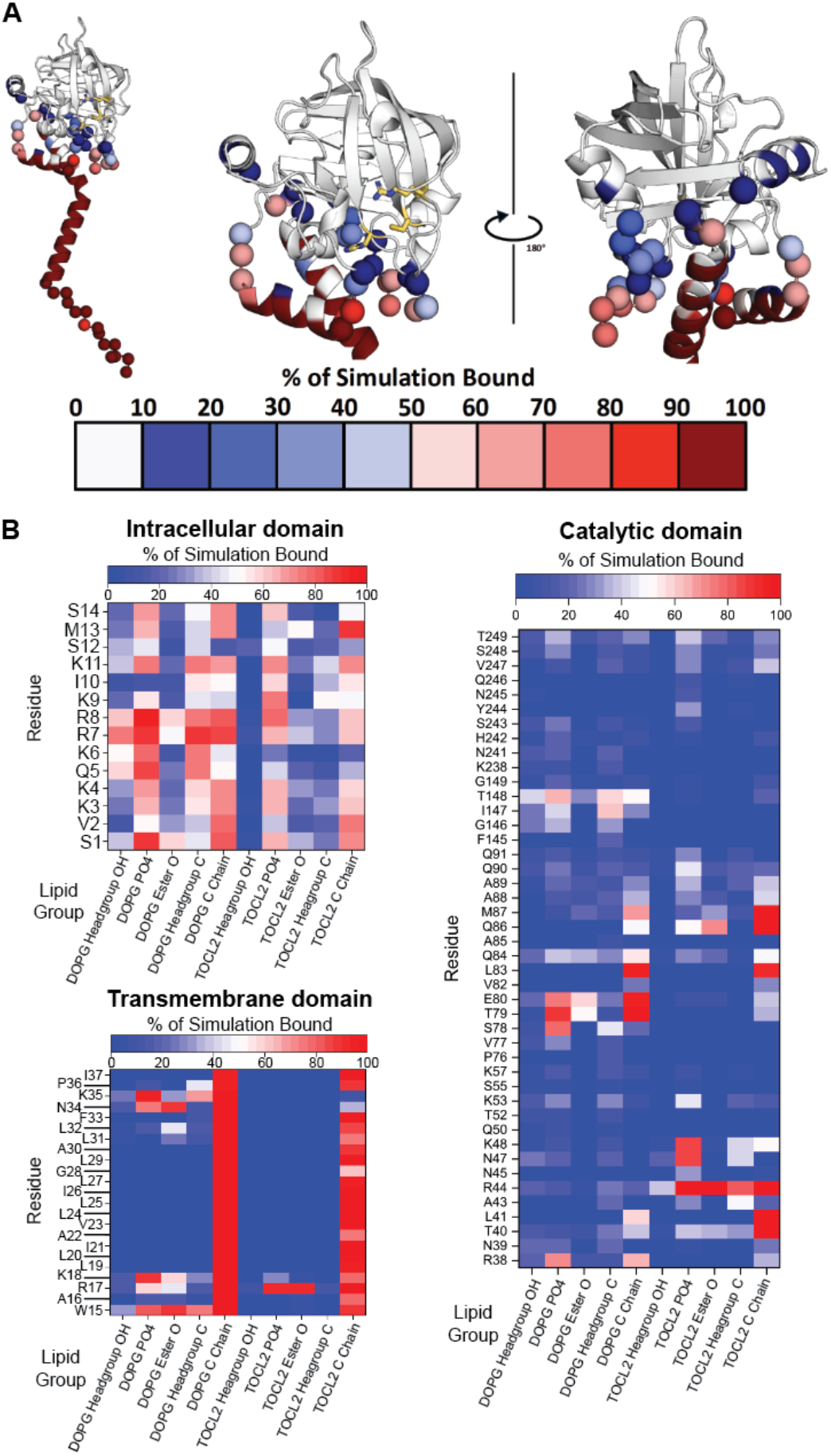
Contact map of spySrtA and the lipid bilayer. Following triplicate 500 ns molecular dynamics simulations, a contact map was generated to assess the percent (%) of simulation bound for specific catalytic domain atoms in spySrtA and lipid groups. (**A**) SpySrtA is shown in cartoon representation and colored in gray. The traditional His-Cys-Arg catalytic residues are in gold, with side chain atoms shown as sticks and colored by heteroatom (N=blue, O=red, C=gold). For amino acids that made specific contacts, the Cα atoms are shown as spheres and colored according to the key. These colors match the data in (**B**). (**B**) Specific contacts for the intracellular, transmembrane, and catalytic domain residues of spySrtA with lipid groups are shown and colored as labeled.

### The transmembrane domains of SpySrtA and its endogenous substrate M protein interact specifically and stably during molecular dynamics simulations

To investigate the tripartite complex of spySrtA, membrane, and substrate, we created two separate models using the pipeline described above. For both, we chose to include the sortase recognition motif initially bound in the active site, in order to investigate interactions between proteins and with the membrane in the bound complex. In future experiments, it would also be interesting to investigate initial recognition of sortase for its substrate(s). In the first model, we used a peptide substrate (LPSTG, where L=P4, P=P3, S=P2, T=P1, and G=P1’) to match the canonical pentapeptide recognition motif (LPXTG) for sortase enzymes (**Figure 3A**).^2,5,6,10^ In addition, this sequence is derived from the *S. pyogenes* M protein, a virulence factor that is attached to the bacterial cell surface by sortase-mediated ligation.^3,60^ Our second model included a region of the M protein containing both the LPSTG sequence and its C-terminal transmembrane domain. Because there is reported variability in M protein extracellular sequences, the protein is very large, and there is no evidence to our knowledge of interactions between the spySrtA catalytic domain and other regions of the substrate, we restricted our model to M protein residues 376-415, which included five residues before the start of the target LPSTG motif (**Figure 3B**). The prediction algorithm TMHMM-2.0 was used to predict the transmembrane domain of M protein, suggesting this domain is residues 387-409.^61^ Our model was largely consistent with this result, although it suggested a more accurate transmembrane domain excludes T387 (**Figure 3C**). For spySrtA, TMHMM-2.0 predicts the transmembrane domain to contain residues 13-32. Again, our model largely agreed, with the addition of F33 and N34 (which interacted with the polar head groups of the lipid molecules) (**Figure 3C**).

**Figure 3.**
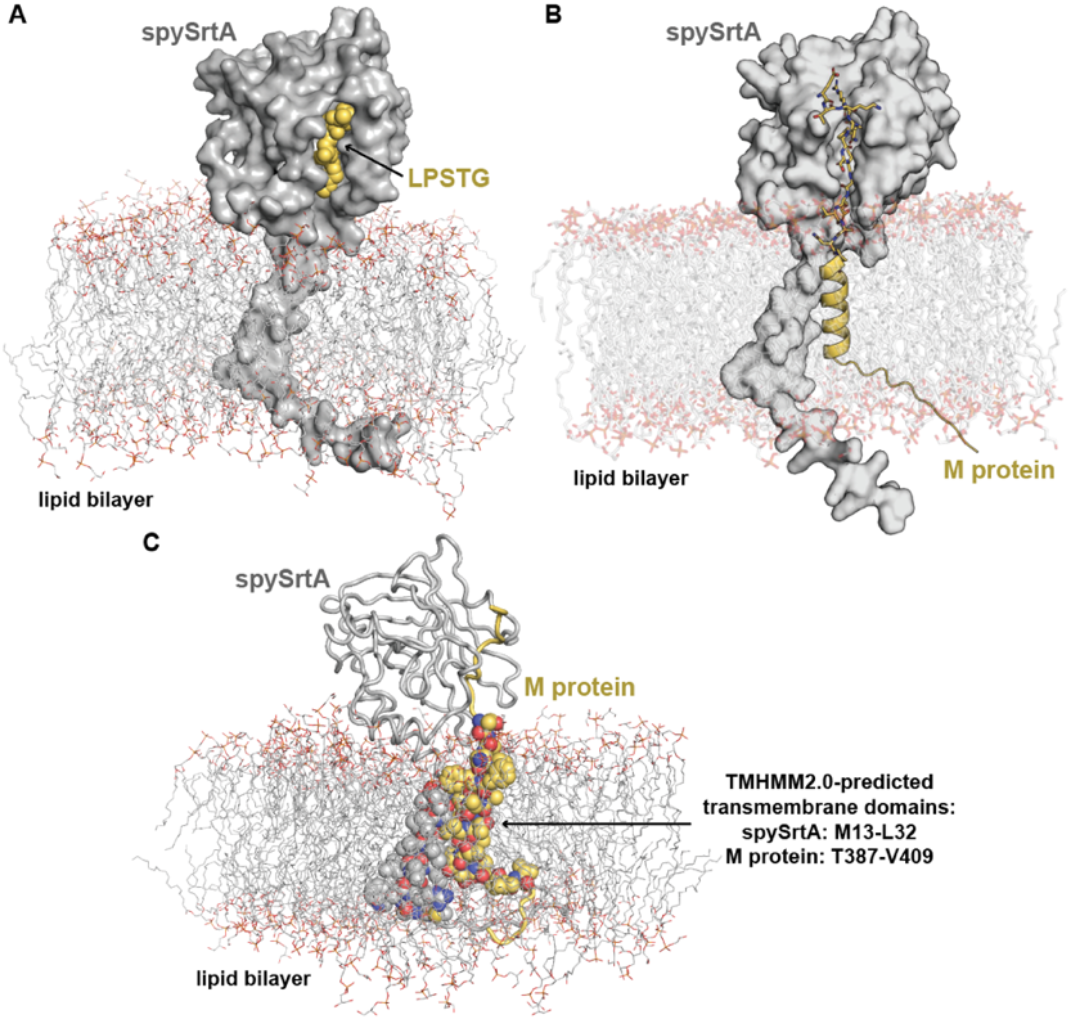
AlphaFold models of full-length spySrtA with substrate inserted into a lipid bilayer. Output models of spySrtA with an LPSTG peptide (**A**) or extended M protein sequence (**B**) are shown with the spySrtA protein in gray surface representation. For all, the lipid bilayer is shown as lines and colored by heteroatom (C=gray, O=red, N=blue). (**A**) The LPSTG peptide is in yellow spheres. (**B**) The extended M protein model is in cartoon for the intracellular and transmembrane domains, and stick representation colored by heteroatom for the extracellular domain (C=yellow), which includes the LPSTG motif. (**C**) The predicted transmembrane domain residues are highlighted as spheres and colored by heteroatom (spySrtA: C=gray, M protein: C=yellow). The extracellular domains are shown in cartoon representation and colored as labeled.

We ran triplicate 500 ns molecular dynamics simulations with each of these spySrtA-substrate-membrane complexes, as described in the Materials and Methods. Again, we observed that all components were stable throughout each simulation, with minimal variability, as measured using relative root-mean-square deviation over time and root-mean-square fluctuation by residue calculations (**Figures S3-4**). This is also apparent in structural alignment of 20 states from an example simulation, with each structure (membrane not shown) representing a state every 25 ns of simulation time (**Figure 4**). The largest variability is seen for the five amino acids preceding the LPSTG motif in M protein, which was not surprising as these residues are not expected to specifically interact with spySrtA (**Figure 4B**). Furthermore, the relative stability of the atoms in the LPSTG sequence in both simulations was similar to what we observed in previous 1000 ns spySrtA_81-249_-LPATA molecular dynamics simulations, using our experimental structures.^26^ The distance between the thiol of the catalytic C208 and P1 carbonyl carbon, the site of nucleophilic attack, was also stabilized by the presence of additional M protein residues, as visualized by a shift in the distance distribution (**Figure S5**). For example, in our triplicate simulations of spySrtA-M protein-membrane (defined as T1, T2, and T3), the distance distribution between these atoms was centered around 3.8 Å for T1 and T2, but closer to 5 Å for T3. For reference, we observed a probability distribution centered at 3.8 Å previously in our spySrtA_81-249_-LPATA simulations.^26^ However, in all three simulations for the spySrtA-LPSTG-membrane system, we saw a bimodal distribution (centered at the 3.8 and 5 Å distances) (**Figure S5**). There is no clear reasoning for this discrepancy, although we predict that this may reflect the peptide and/or substrate sampling both a catalytically competent bound state and an unreactive partially bound state. With M protein, the averaged ratio favors the closer or ‘bound’ state, with relative probabilities of roughly 0.25:0.15 for the peak maxima of 3.8 Å:5 Å, or ‘bound’:’partially bound’ distances. Conversely, for the LPSTG peptide simulations, this ratio is flipped, at roughly 0.2:0.3 for the peaks corresponding to the 3.8 Å:5 Å distances (**Figure S5**). Notably, the peptide does stay stably bound despite some fluctuations in this distance (**Figure 4**).

**Figure 4.**
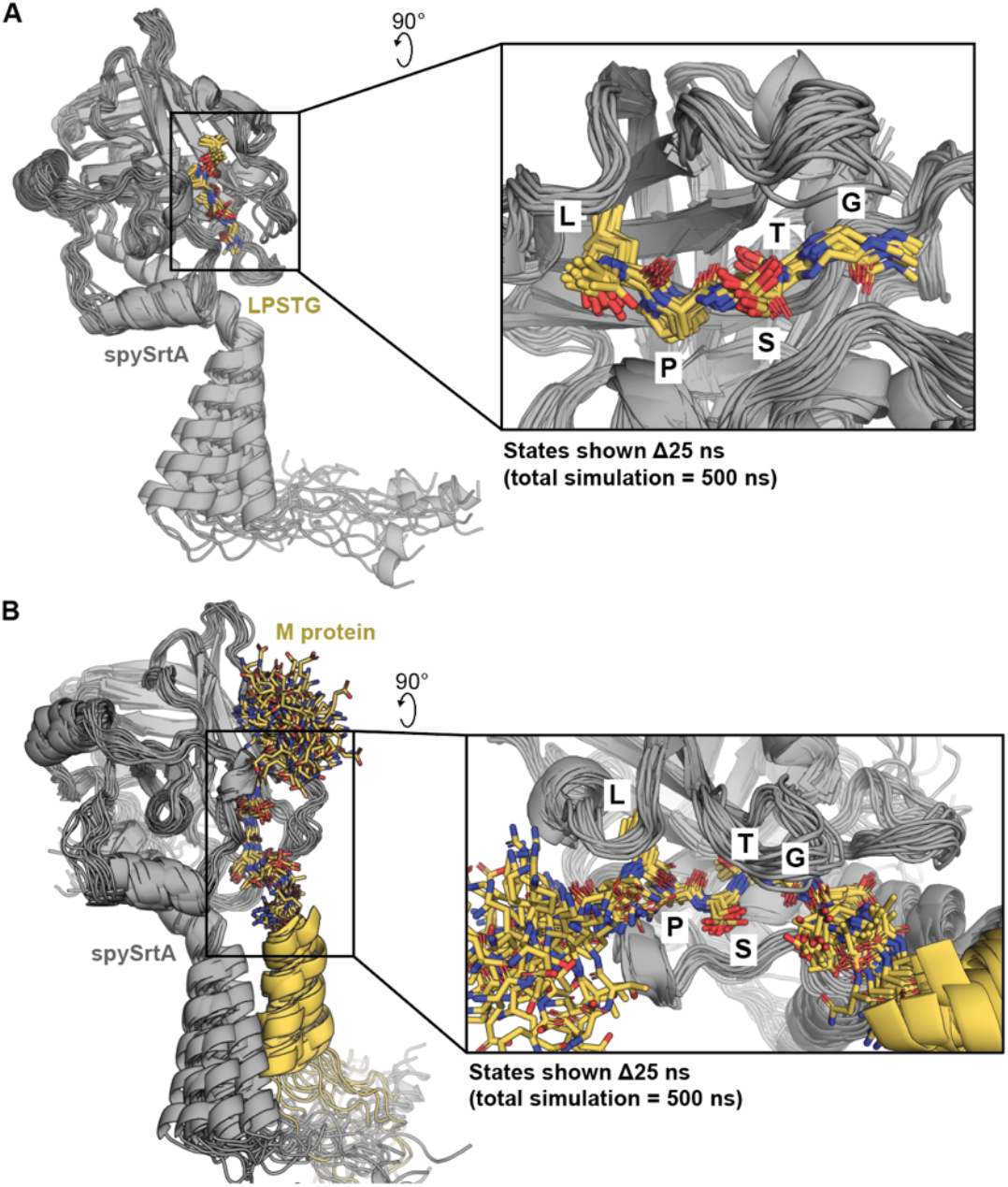
Molecular dynamics simulations reveal stable substrate binding to spySrtA. The results of one simulation replicate are shown for spySrtA-LPSTG (**A**) and spySrtA-M protein (**B**). Output states corresponding to Δt=25 ns (21 states total, including t=0) are aligned and shown. The lipid bilayer is not shown for clarity (although it was present in all simulations). SpySrtA is in gray cartoon. The intracellular and transmembrane domain of M protein is in yellow cartoon (**B**). All other peptide or M protein (LPSTG or KRQLPST, respectively) residues are shown as yellow sticks and colored by heteroatom (N=blue, O=red, C=yellow). The insets show a zoomed-in version of the interaction and are rendered similarly.

The spySrtA-M protein-membrane model and simulations also revealed a relatively stable transmembrane domain interaction between the two proteins. This interaction persists throughout the entirety of each replic ate simulation, although the specific residues that maintain contact varies (**Figure 5**). Here, we visually analyzed amino acids oriented towards each other at t=0 ns of each simulation, including Leu20, Ile21, Leu24, Gly28, and Leu31 in spySrtA and Phe392, Ala395, Ala396, Val399, and Ala403 in M protein, for the t=0, 250, and 500 ns states (**Figure 5**). The transmembrane domains of these proteins remain associated in all replicates, with the highest degree of dissociation at t=500 ns for simulation T1. Furthermore, in the t=500 ns state, interacting residues differ for T1 (Leu31-Ala395), T2 (Leu24-Ala403 and Leu31-Ala396), and T3 (Leu31-Phe392, Leu24-Val399, and Ile21-Ala403) (**Figure 5**). Overall, these simulations suggest that the transmembrane domain interaction between spySrtA and M protein may be multivalent, likely reflecting the dynamic nature of both the membrane and proteins themselves.

**Figure 5.**
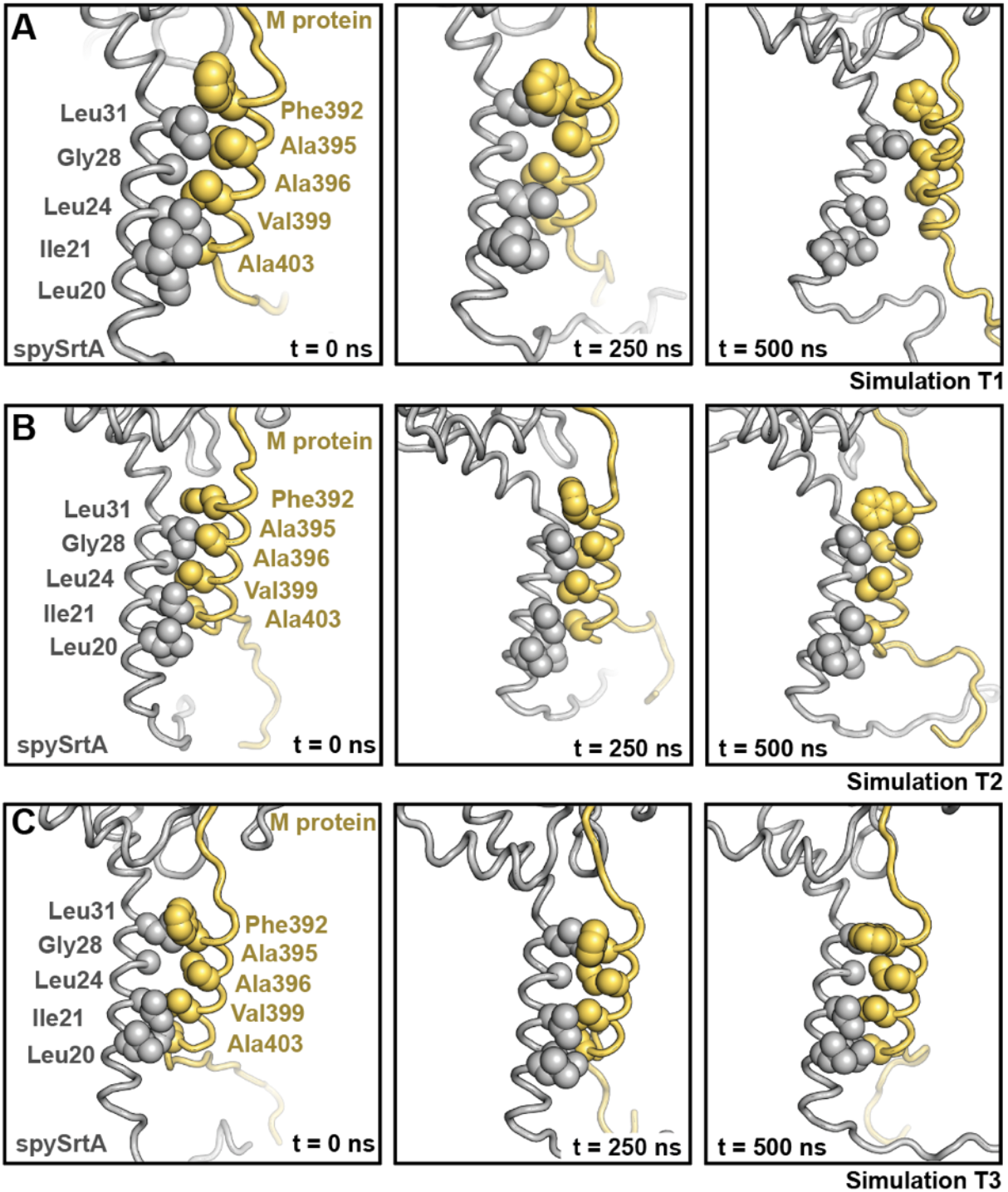
Structural trajectories of spySrtA and M protein transmembrane domain interactions. Specific states (corresponding to t=0, 250, 500 ns) are shown for each replicate simulation, T1 (**A**), T2 (**B**), and T3 (**C**). The lipid bilayer is not shown for clarity although is present in all simulations. SpySrtA is shown as gray cartoon and M protein as yellow cartoon. Amino acid sidechains that are oriented towards each other in the transmembrane helices are shown as spheres and labeled for all.

### Specific residues in spySrtA and M protein are buried upon substrate binding

We wanted to further analyze substrate recognition by the catalytic domain of spySrtA in the context of the full-length proteins. When we analyzed the change in solvent accessible surface area (ΔSASA), defined as SASA_bound_-SASA_unbound_ (in Å^2^), we see that the P4 Leu and P1 Thr in the M protein fragment are substantially buried (−ΔSASA > 10 Å^2^) upon substrate binding in the triplicate simulations (**Figures 6A-B**). In addition, we saw that for one of the replicates, the largest -ΔSASA value observed was for the P2’ Glu. For the other two replicates, -ΔSASA for the P2’ Glu was second to only the P4 Leu, a position previously described as binding in a specific hydrophobic pocket (**Figure 6B**).^26^ This observation will be discussed in detail below. Other substrate residues with relatively large -ΔSASA values include the P6 Arg (**R**Q<u>LPSTG</u>E,

**Figure 6.**
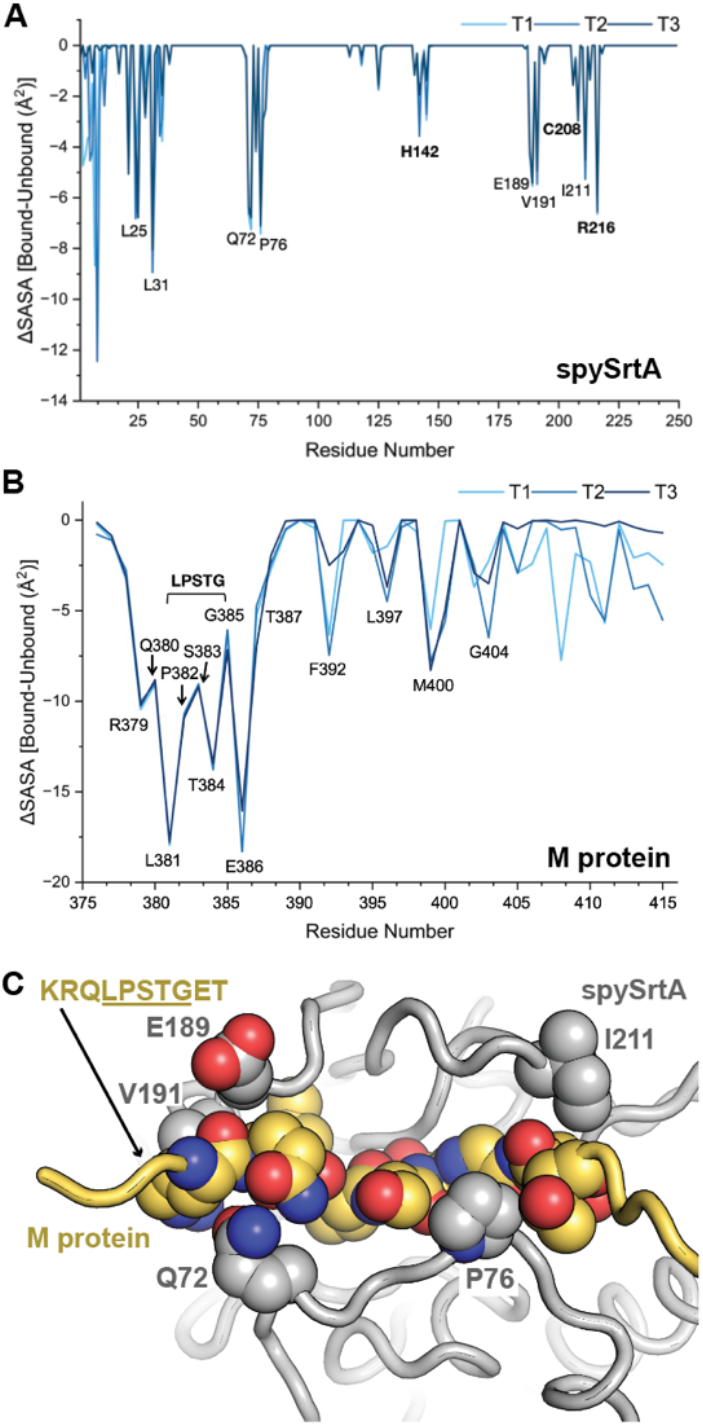
Residues outside the canonical pentapeptide recognition motif interact with the catalytic domain of spySrtA. Analysis of the change in solvent accessible surface area (SASA) between the bound and unbound AlphaFold models (ΔSASA) reveal several amino acids that become buried upon substrate binding, defined as a relatively large -ΔSASA are highlighted and labeled, for spySrtA (**A**) and M protein (**B**). (**C**) Predicted interactions at amino acids N-terminal (KRQ) and C-terminal (ET) to the LPSTG pentapeptide recognition motif are highlighted in spySrtA (gray cartoon) as side chain spheres and colored by heteroatom (C=gray, O=red, N=blue). The KRQLPSTGET sequence of M protein is shown as spheres and colored by heteroatom (C=yellow), with other amino acids as a yellow cartoon. M protein numbering is based on the full-length protein.

Arg in **bold**), P3 Pro, P2 Ser, P1’ Gly, and other positions within the transmembrane domain (**Figure 6B**).

For the spySrtA enzyme, in addition to the expected positions required for enzymatic activity (H142-C208-R216), we also observed other residues within the catalytic domain with a similar -ΔSASA, including E189, V191, and I211 (**Figure 6A**). These positions may bind and stabilize substrate binding outside the LPSTG recognition motif, for example at the N-terminal KRQ and C-terminal ET positions (M protein sequence = KRQ<u>LPSTG</u>ET) (**Figure 6C**). Taken together, these results suggested there may be specific interactions between spySrtA and M protein beyond the canonical LPXTG recognition motif for class A sortases.

Two additional spySrtA residues (Q72 and P76) that are not in the transmembrane domain also exhibited relatively large -ΔSASA values in the substrate bound models (**Figure 6A**). These two residues also appear to interact directly with the M protein substrate in or near the recognition motif, at either the P5 Gln (with Q72) or the P2 Ser and P1’ Gly (with P76) positions (**Figure 6C**). These interactions are present in all three replicate simulations, with average distances between Cα atoms equal to: Q72-P5 Gln = 7.3 ± 0.5 Å (T1), 7.4 ± 0.6 Å (T2), and 7.3 ± 0.5 Å (T3), and P76-P1’ Gly = 6.1 ± 0.7 Å (T1), 4.8 ± 0.3 Å (T2), and 6.3 ± 0.7 Å (T3).

### The extracellular domain (amino acids 34-249) of spySrtA interacts with its endogenous substrate M protein at positions beyond the canonical pentapeptide recognition motif

To complement the analysis of our AlphaFold models, we also used our spySrtA_1-249_-M protein_376-415_ molecular dynamics simulations to investigate interactions in residues adjacent to the LPXTG substrate recognition motif. In our triplicate simulations, contact map analysis revealed interactions at amino acids both N- and C-terminal to the M protein LPSTG sequence (**Figure 7A**). At the N-terminus, these data confirmed interactions between Q72, E189 and V191 with the P6-P5 RQ (<u>RQ</u>LPSTG) positions, as also highlighted above in our ΔSASA analysis. A potential role for backbone atoms of F71 and P188 was also identified (**Figure 7A**). We observed even more persistent interactions adjacent to the C-terminus of LPSTG, with both specific backbone and side-chain contacts between the P2’ Glu (LPSTG<u>E</u>) and several spySrtA residues. Most notably, electrostatic interactions with the side-chain atoms of K35 and R38 in spySrtA were present throughout each simulation (**Figure 7B**).

**Figure 7.**
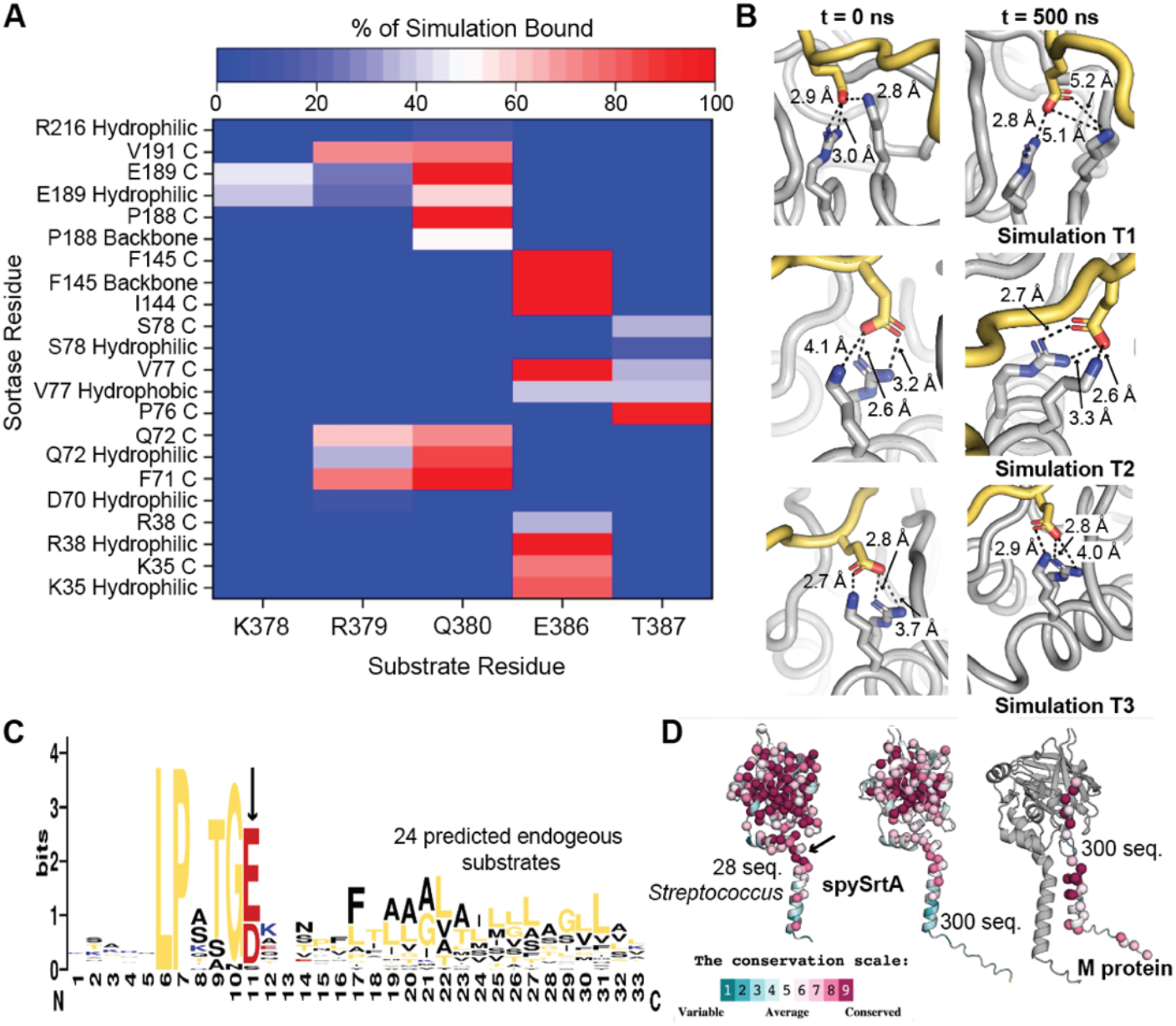
Additional specific interactions are identified between spySrtA and the P2’ Glu in M protein. (**A**) Contact map of spySrtA amino acids and either the KRQ or ET residues of M protein, in the sequence KRQLPSTGET. “Hydrophilic” refers to side chain atoms. (**B**) Initial and final states from the T1, T2, and T3 simulations (t=0 and 500 ns) highlighting persistent interactions between K35 and R38 spySrtA with the P2’ Glu in M protein. M protein is shown as yellow cartoon with the P2’ Glu side chain as sticks and colored by heteroatom (C=yellow, O=red, N=blue). SpySrtA is shown as gray cartoon with the K35 and R38 side chain atoms as sticks and colored by heteroatom (C=gray). Relevant distances are labeled. (**C**) Sequence logo (WebLogo) of 24 predicted endogenous substrates of spySrtA confirms conservation at the P2’ position for a negatively-charged amino acid (either D or E). (**D**) ConSurf analysis with 28 *Streptococcus* SrtA sequnces reveals that the K35 and R38 positions are generally conserved (left), although this is not true for 300 SrtA sequences from a broader range of bacterial species (middle). M protein conservation is also highlighted (right). For all, the proteins are shown in cartoon representation with relevant Cα atoms as spheres. The conservation scale is shown and labeled.

Multiple sequence alignment of spySrtA plus 27 additional *Streptococcus* SrtA proteins indicates that these Lys and Arg positions are relatively well conserved.

Nineteen of the 28 sequences contain a Lys in the equivalent K35 position, with the other sequences containing either Ser or Thr polar residues (**Figure S6**). Twenty-five of the 28 sequences contain an Arg in the equivalent R38 position (**Figure S6**). Conservation is also reflected in spySrtA substrate sequences; a sequence logo of 24 predicted spySrtA endogenous substrates reveals that a negative amino acid (either Glu or Asp) in the P2’ position is highly conserved (**Figure 7C**).

We also used Consurf to investigate the evolutionary conservation of these residues visually (**Figure 7D**).^55,56^ When we limited our analysis to the 28 *Streptococcus* SrtA proteins, again, the relatively high conservation of residues, e.g., K35 and R38 in spySrtA, were apparent (black arrow in left panel of **Figure 7D**); however, when applied to 300 SrtA sequences (middle panel in **Figure 7D**), this was not conserved. For example, sequence alignment of spySrtA with *Staphylococcus aureus* SrtA (UniProt ID SRTA_STAA8) revealed that while the Lys is conserved (K26 in *S. aureus* SrtA), the residue in the equivalent Arg position is D30. To our knowledge, these types of interactions have not been explored in the literature, and it remains unclear whether they have a significant impact on spySrtA activity. In the first step of sortase-mediated catalysis, the substrate is cleaved between the P1/P1’ positions (LPST/GE), and initial binding of the substrate is potentially facilitated by interactions outside of the standard LPXTG motif.^2,9,10^ Additional experimentation will be necessary to probe these interactions further, and to understand the mechanistic implications of these observations.

## Discussion

One of our major questions prior to modeling full-length spySrtA with M protein in the membrane was to understand how the LPXTG recognition motif would be oriented with respect to the enzyme active site. The transmembrane domain of M protein is C-terminal to the LPSTG by only a small number of residues, and the stereochemistry of recognition is unclear. A transmembrane domain prediction algorithm predicted it starts at T387, just two amino acids after the P1’ Gly, sequence: ^381^<u>LPSTG</u>ETANPFF^392^. Our model was able to illustrate that the enzyme and substrate are indeed well positioned for catalysis in the context of a lipid bilayer, which, while not surprising from an evolutionary or biochemical standpoint, provides a new structural view of this fundamentally important enzymatic reaction. In addition, we observed that because the sortase recognition motif is proximal to the membrane, the catalytic domain of spySrtA was also positioned near the membrane surface, and our contact map revealed several residues in the catalytic domain that interacted directly with lipid molecules (**Figure 1C**).

Another insight that resulted from our simulations was that there were interactions between spySrtA and M protein residues outside of the LPSTG pentapeptide recognition motif. This included the following amino acids (underlined), <u>KRQ</u>LPSTG<u>ET</u> (**Figure 7A**). Most notably, we observed specific interactions between K35 and R38 in spySrtA, which are well conserved in *Streptococcus* SrtA enzymes, and the P2’ Glu, which is also strongly conserved in endogenous substrates for spySrtA (**Figures 7C, S6**). The contribution of these contacts was not tested with respect to the sortase-mediated ligation reaction, however, we predict that a better understanding of specificity at these positions may be useful in the design of substrates for protein engineering applications using sortase-mediated ligation (SML) strategies.^10,14^ Consistent with these observations, our data also suggested that the presence of the transmembrane domain of M protein may facilitate an increased percentage of ‘bound’ substrate in the active site as compared to the isolated peptide (**Figure S5**).

This is a challenging system to study experimentally. In preliminary experiments not presented here, we successfully purified full-length spySrtA in lipid nanodiscs and confirmed catalytic activity with a fluorescent peptide substrate, similar to our previous work.^26,62–64^ Significantly, the ideal substrate would also include its transmembrane domain in order to directly probe the role of the transmembrane domain in substrate recognition, as well as to test a potential transmembrane domain-mediated interaction between spySrtA with M protein. Challenges in the system design included controlling the membrane insertion direction for both enzyme and substrate, and inhibiting enzymatic activity prior to experiment initiation. These issues are surmountable, and experimental work is ongoing.

Overall, our molecular dynamics simulations revealed structural insights into the substrate recognition of a class A sortase with an endogenous substrate within a model lipid bilayer. Specifically, we identified specific contacts between the enzyme and membrane, and our results indicated that there is a transmembrane domain interaction that may facilitate sortase catalysis *in vivo*. Specifically, we hypothesize that the proposed transmembrane interaction could enhance both substrate recognition and turnover rate *in vivo*, thus directly impacting relative enzyme efficiency. This has potential implications for SML protein engineering methods, where the catalytic domains of class A sortases in isolation are widely used despite the wild-type enzymes being limited by low catalytic efficiencies.^14,15,65,66^ While gains in enzyme efficiency and substrate scope have been achieved via directed evolution experiments and other strategies, we envision that a complementary approach may be to mimic the novel enzyme-substrate interactions described here.^13,15,16,67,68^ In this regard, prior work has demonstrated that preassociation of sortase and sortase substrates through either protein-protein interactions or on the surface of liposomes does indeed facilitate SML.^69–71^ The continued development of these strategies is thus a promising means for the further development of SML methodology.

## Acknowledgements

We want to thank all additional members of the Amacher, Antos, McCarty, and Spiegel labs for helpful discussion and research support. This work was supported by NIH 1R15GM154315-01 to J.F. Amacher, J.M. Antos, and J. McCarty. It was additionally supported by NSF CHE-2044958 and a Cottrell Scholar Award from the Research Corporation for Science Advancement to J.F. Amacher, NSF CHE-2102189 and MCB-2441210 to J. McCarty, and NIH 2R15HL135658-03 to P.C. Spiegel.

