## Supporting Information for "An interaction between the transmembrane domains of *Streptococcus pyogenes* sortase A and its endogenous substrate M protein revealed by molecular dynamics simulations"

**Table of Contents:**

**Table S1. Details of the molecular dynamics simulation size.** 2

**Figure S1. Prediction of transmembrane regions in spySrtA and M protein.** 3

**Figure S2. RMSF and RMSD values of triplicate molecular dynamics simulations for full-length spySrtA in a lipid bilayer.** 4

**Figure S3. RMSF and RMSD values of triplicate molecular dynamics simulations for full-length spySrtA with the LPSTG peptide in a lipid bilayer.** 5

**Figure S4. RMSF and RMSD values of triplicate molecular dynamics simulations for full-length spySrtA with M protein in a lipid bilayer, and averaged RMSF values.** 6

**Figure S5. Distribution of distances between the C208 thiol and P1 Thr carbonyl carbon for all simulations.** 7

**Figure S6. Multiple sequence alignment of 28 *Streptococcus* SrtA sequences.** 8-9

**Sequences used for AlphaFold2 modeling.** 10

**Table S1. Details of the molecular dynamics simulation size.**

| System | Total number of atoms | Cubic box dimensions [nm] | Simulation time [ns], N=3 (T1,T2,T3) |
|---|---|---|---|
| SpySrtA (Apo) | 89742 | 8.12781 x 8.12781 x 13.25244 | 500 |
| SpySrtA-LPSTG | 87925 | 8.04625 x 8.04625 x 13.20473 | 500 |
| SpySrtA-M Protein | 91827 | 8.18071 x 8.18071 x 13.41281 | 500 |

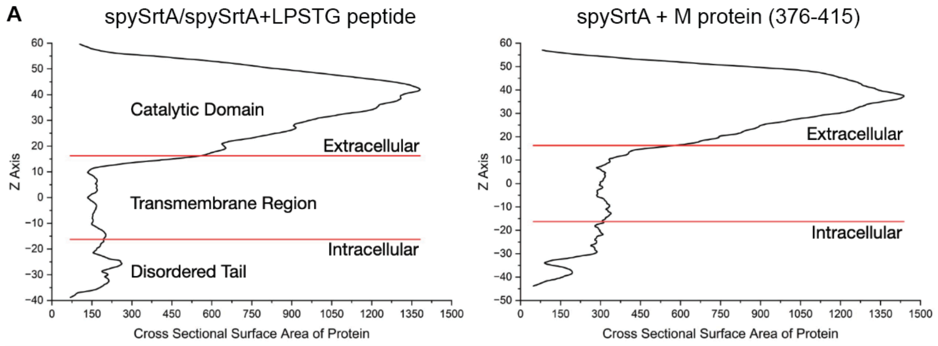


B

| Construct | Depth/Hydrophobic Thickness (Å) | $\Delta G_{transfer}$ (kcal/mol) | Tilt Angle | Membrane Embedded Residues |
|---|---|---|---|---|
| spySrtA | 30.2 ± 1.4 | -28.7 kcal/mol | 21.± 3.° | 13-37,83 |
| spySrtA and substrate | 27.0 ± 1.9 | -35.5 kcal/mol | 18.± 1.° | 14-34, 36-37, 47-48 (spySrtA)<br><br>384-402, 404 (M protein) |

**Figure S1. Prediction of transmembrane regions in spySrtA and M protein.** PPM2.0 was used to predict the transmembrane domain boundaries for spySrtA in output AlphaFold2 models that either contained just full-length spySrtA or spySrtA plus an LPSTG peptide or extended M protein sequence, as labeled. (**A**) The cross sectional surface area of the protein is graphed as a function of the Z Axis, or distance from center of the lipid bilayer (defined as equal to 0). The globular extracellular domain is predicted to have a larger cross sectional surface area than either the transmembrane domain or disordered tail. **(B)** Details of membrane insertion of spySrtA with and without substrate.

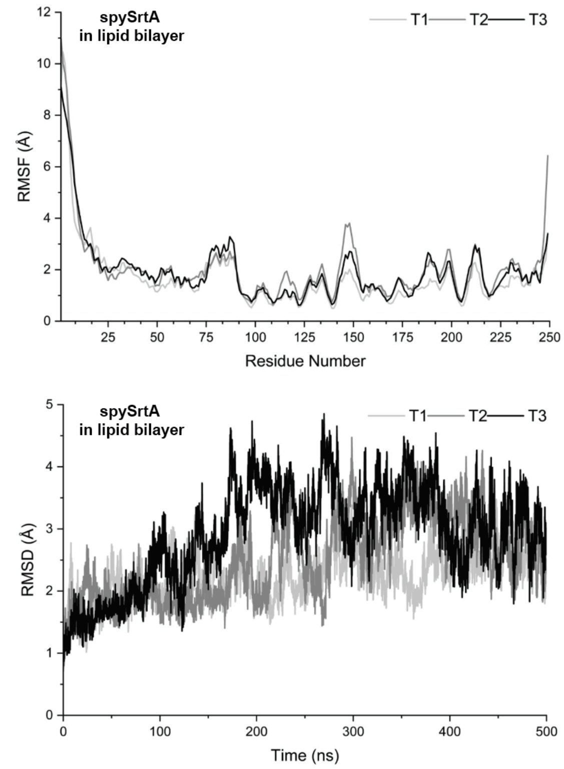


**Figure S2. RMSF and RMSD values of triplicate molecular dynamics simulations for full-length spySrtA in a lipid bilayer.** The root-mean-square-fluctuation (RMSF), or average displacement from their average position, for each residue (using the C$\alpha$ atoms) in full-length spySrtA is shown for each simulation replicate: T1, T2, and T3 (**top**). This highlights regions of the protein with relatively higher degrees of flexibility. The room-mean-square-deviation (RMSD) for the averaged spySrtA protein over each 500 ns simulation is shown for each simulation replicate: T1, T2, and T3 (**bottom**).

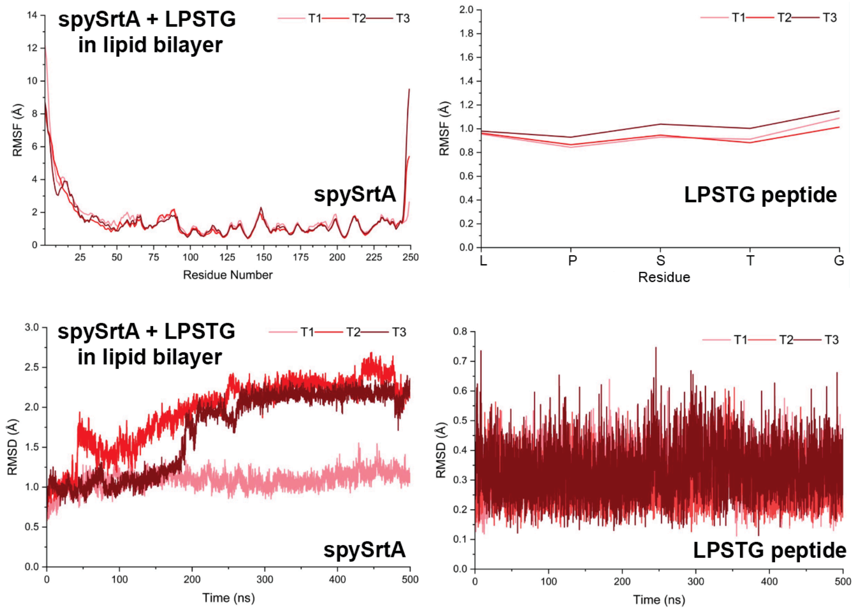


**Figure S3. RMSF and RMSD values of triplicate molecular dynamics simulations for full-length spySrtA with the LPSTG peptide in a lipid bilayer.** The root-mean-square-fluctuation (RMSF), or average displacement from their average position, for each residue (using the Cα atoms) in full-length spySrtA (**left**) or the LPSTG peptide (**right**) is shown for each simulation replicate: T1, T2, and T3 (**top**). The room-mean-square-deviation (RMSD) for the averaged spySrtA protein (**left**) or LPSTG peptide (**right**) over each 500 ns simulation is shown for each simulation replicate: T1, T2, and T3 (**bottom**).

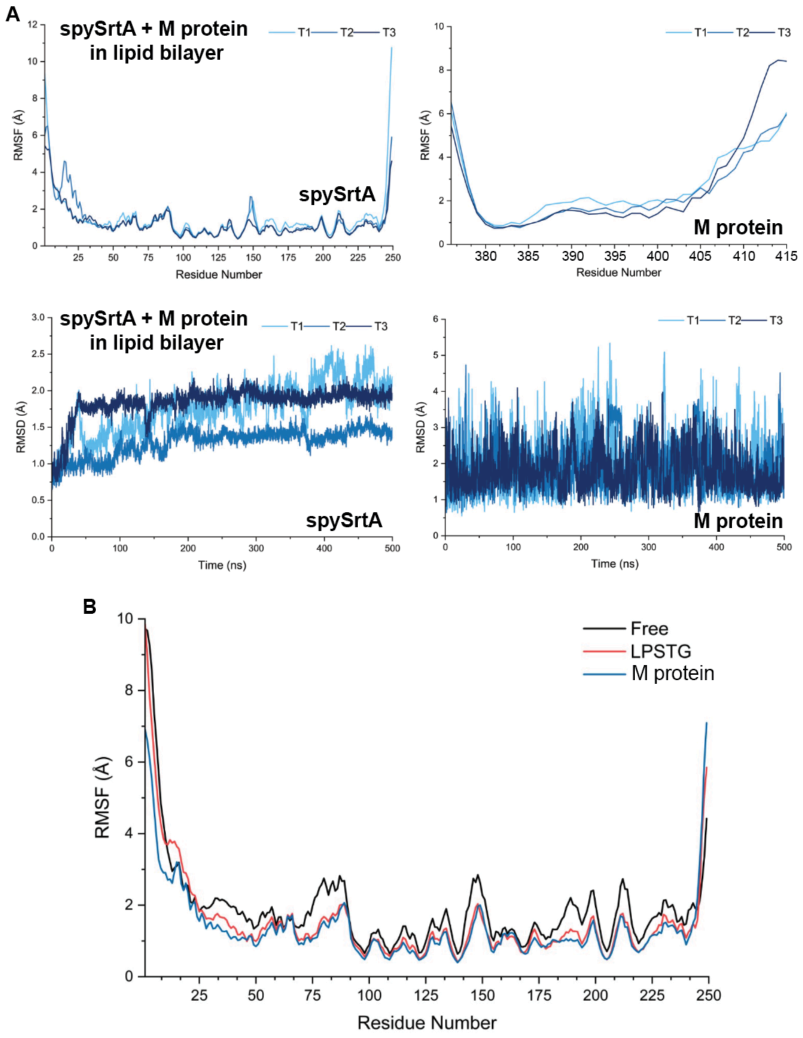


**Figure S4. RMSF and RMSD values of triplicate molecular dynamics simulations for full-length spySrtA with M protein in a lipid bilayer, and averaged RMSF values.** (**A**) The root-mean-square-fluctuation (RMSF), or average displacement from their average position, for each residue (using the C$\alpha$ atoms) in full-length spySrtA (**left**) or M protein (**right**) is shown for each simulation replicate: T1, T2, and T3 (**top**). The room-mean-square-deviation (RMSD) for the averaged spySrtA protein (**left**) or M protein (**right**) over each 500 ns simulation is shown for each simulation replicate: T1, T2, and T3 (**bottom**). (**B**) Averaged RMSF values for spySrtA in the full-length spySrtA ("free"), spySrtA+LPSTG ("LPSTG"), and spySrtA+M protein ("M protein") triplicate simulations is shown.

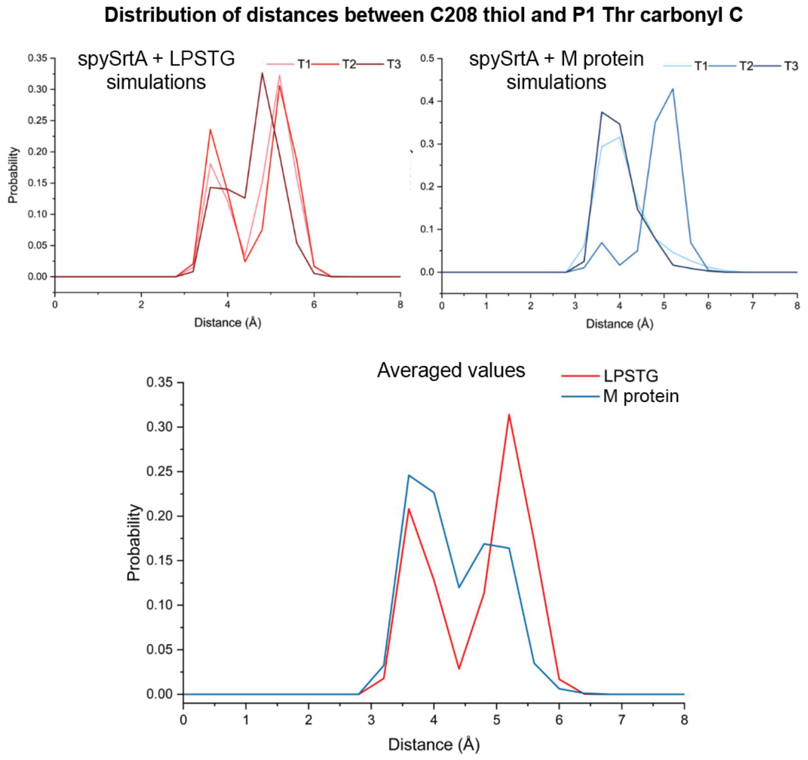


**Figure S5. Distribution of distances between the C208 thiol and P1 Thr carbonyl carbon for all simulations.** The distance for each state of a replicate simulation (T1, T2, and T3) between the C208 thiol and its site of nucleophilic attack, the P1 Thr carbonyl carbon, was graphed as a distribution and is shown for the spySrtA+LPSTG peptide simulations (**top left graph**) and spySrtA+M protein simulations (**top right graph**). Averaged values over triplicate simulations are shown in the **bottom graph**, as labeled (spySrtA+LPSTG = red, spySrtA+M protein = blue).

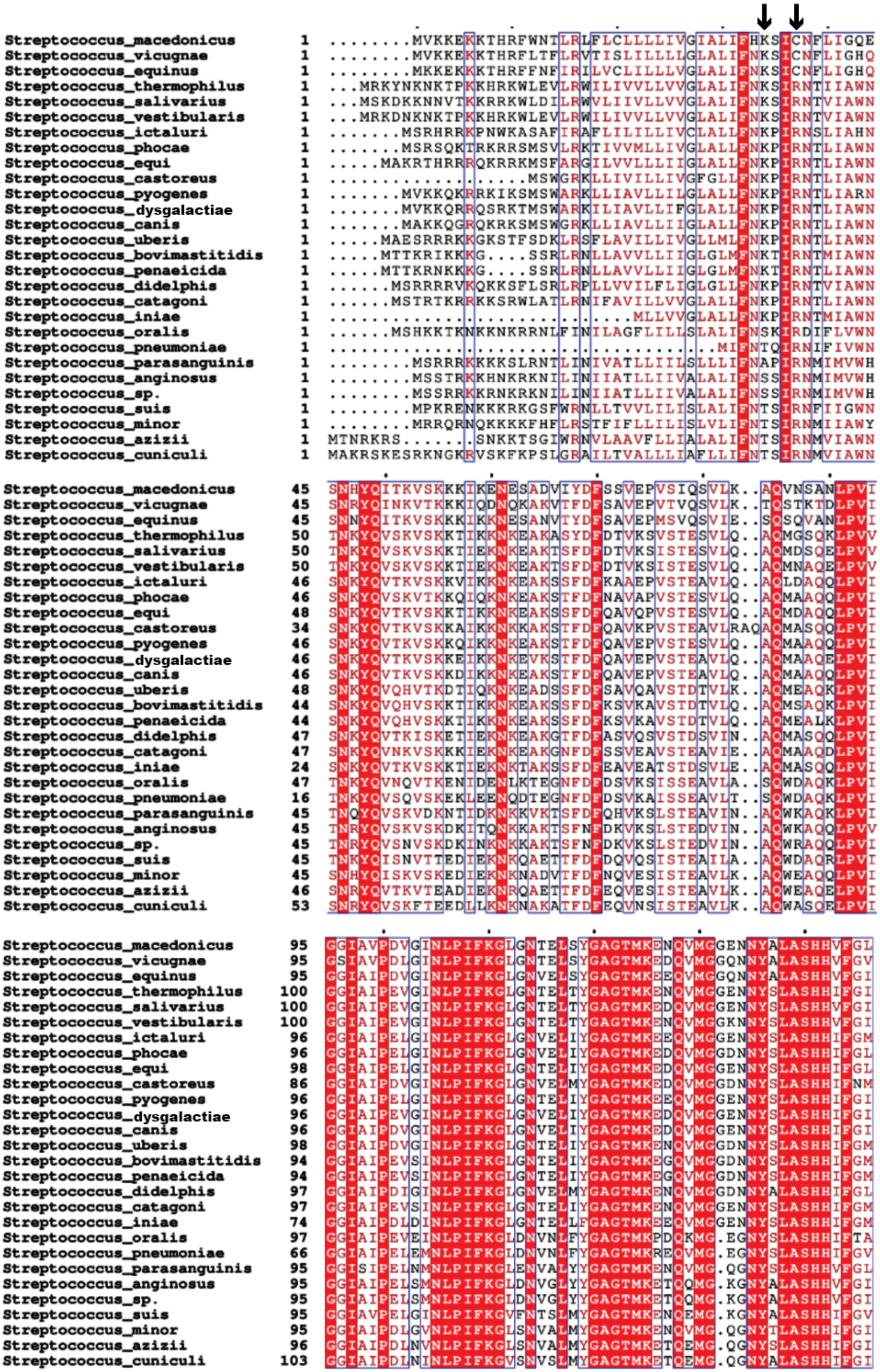
Streptococcus_macedonicus 1 ........MVKKEKKTHRFWNTLRLFLCLLLLIVGIALIFHKSICNFLIGQE
Streptococcus_vicugnae 1 ........MVKKEKKTHRFLTFLRVTISLILLLLGLALIFNKSICNFLIGHQ
Streptococcus_equinus 1 ........MKKEKKKTHRFFNFIRILVCLILLLVGLALIFNKSICNFLIGHQ
Streptococcus_thermophilus 1 ...MRKYNKNKTPKKHRKWLEVLRWILIVVLLVVGLALIFNKSIRNTIIAWN
Streptococcus_salivarius 1 ...MSKDKKNNVTKKRRKWLDILRWVLIIVLLVVGLALIFNKSIRNTVIAWN
Streptococcus_vestibularis 1 ...MRKDNKNKTPKKHRKWLEVLRWILIVVLLVVGLALIFNKSIRNTVIAWN
Streptococcus_ictaluri 1 .......MSRHRRKPNWKASAFIRAFLILILLIVCLALLFNKPIRNSLIAHN
Streptococcus_phocae 1 .......MSRSQKTRKRRSMSVLRKTIVVMLLIVGLALLFNKPIRNTLIAWN
Streptococcus_equi 1 .....MAKRTHRRRQKRRKMSFARGILVVLLLIIGLALLFNKPIRNTLIAWN
Streptococcus_castoreus 1 ...................MSWGRKLLIVILLIVGFGLLFNKPIRNTLIAWN
Streptococcus_pyogenes 1 .......MVKKQKRRKIKSMSWARKLLIAVLLILGLALLFNKPIRNTLIARN
Streptococcus_dysgalactiae 1 .......MVKKQRRQSRKTMSWARKILIAVLLIFGLALLFNKPIRNTLIAWN
Streptococcus_canis 1 .......MAKKQGRQKRKSMSWGRKLLIAVLLIIGLALLFNKPIRNTLIAWN
Streptococcus_uberis 1 .....MAESRRRKKGKSTFSDKLRSFLAVILLIVGLLMLFNKPIRNTLIAWN
Streptococcus_bovimastitidis 1 .....MTTKRIKKKG....SSRLRNLLAVVLLIIGLGLMFNKTIRNTMIAWN
Streptococcus_penaeicida 1 .....MTTKRNKKKG....SSRLRNLLAVVLLIIGLGLMFNKTIRNTLIAWN
Streptococcus_didelphis 1 ......MSRRRRVKQKKSFLSRLRPLLVVILFLIGLGLLFNKPIRNTVIAWN
Streptococcus_catagoni 1 ......MSTRTKRRKKSRWLATLRNIFAVILLVVGLALLFNKPIRNTLIAWN
Streptococcus_iniae 1 .............................MLLVVGLALLFNKPIRNTMIAWN
Streptococcus_oralis 1 ......MSHKKTKNKKNKRRNLFINILAGFLILLSLALIFNSKIRDIFLVWN
Streptococcus_pneumoniae 1 .....................................MIFNTQIRNIFIVWN
Streptococcus_parasanguinis 1 ........MSRRRKKKKSLRNTLINIVATLLIILSLLLIFNAPIRNMIMVWH
Streptococcus_anginosus 1 ........MSSTRKKHNKRKNILINIIATLLIIVALALIFNSSIRNMIMVWH
Streptococcus_sp. 1 ........MSSRRKKRNKRKNILINIIATLLIIVALALIFNSSIRNMIMVWH
Streptococcus_suis 1 ........MPKRENKKKRKGSFWRNLLTVVLILISLALIFNTSIRNFIIGWN
Streptococcus_minor 1 ........MRRQRNQKKKKFHFLRSTFIFLLILISLVLIFNTSIRNMIIAWY
Streptococcus_azizii 1 MTNRKRS.......SNKKTSGIWRNVLAAVFLLIALALIFNTSIRNMVIAWN
Streptococcus_cuniculi 1 MAKRSKESRKNGKRVSKFKPSLGRAILTVALLLIAFLLIFNTSIRNMVIAWN
Streptococcus_macedonicus 45 SNHYQITKVSKKKIKENESADVIYDFSSVEPVSIQSVLK..AQVNSANLPVI
Streptococcus_vicugnae 45 SNRYQINKVTKKKIQDNQKAKVTFDFSAVEPVTVQSVLK..TQSTKTDLPVI
Streptococcus_equinus 45 SNNYQITKVSKKKIKKNESANVTYDFSAVEPMSVQSVIE..SQSQVANLPVI
Streptococcus_thermophilus 50 TNKYQVSKVSKKTIKKNKEAKASYDFDTVKSVSTESVLQ..AQMGSQKLPVV
Streptococcus_salivarius 50 TNKYQVSKVSKKTIEKNKEAKTSFDFDTVKSISTESVLQ..AQMDSQELPVV
Streptococcus_vestibularis 50 TNKYQVSKVSKKTIEKNKEAKTSFDFDTVKSISTESVLQ..AQMNAQELPVV
Streptococcus_ictaluri 46 SNKYQVTKVSKKVIKKNKEAKSSFDFKAAEPVSTEAVLQ..AQLDAQQLPVI
Streptococcus_phocae 46 SNKYQVSKVTKKQIQKNKEAKSTFDFNAVAPVSTESVLQ..AQMAAQQLPVI
Streptococcus_equi 48 SNKYQVTKVSKKTIKKNKEAKSSFDFQAVQPVSTESVLQ..AQMDAQQLPVI
Streptococcus_castoreus 34 SNKYQVTKVSKKAIKKNKKAKSTFDFQAVKPVSTEAVLRAQAQMASQQLPVI
Streptococcus_pyogenes 46 SNKYQVTKVSKKQIKKNKEAKSTFDFQAVEPVSTESVLQ..AQMAAQQLPVI
Streptococcus_dysgalactiae 46 SNKYQVTKVSKKEIKKNKEVKSTFDFQAVEPVSTEAVLQ..AQMAAQELPVI
Streptococcus_canis 46 SNKYQVTKVSKKDIKKNKEAKSTFDFQAVEPVSTEAVLQ..AQMAAQQLPVI
Streptococcus_uberis 48 SNKYQVQHVTKDTIQKNKEADSSFDFSAVQAVSTDTVLK..AQMEAQKLPVI
Streptococcus_bovimastitidis 44 SNKYQVQHVSKKTIKKNKEAKSSFDFKSVKAVSTDTVLQ..AQMSAQKLPVI
Streptococcus_penaeicida 44 SNKYQVQHVSKKTIKKNKEAKSSFDFKSVKAVSTDTVLQ..AQMEALKLPVI
Streptococcus_didelphis 47 SNKYQVTKISKETIEKNKEAKGTFDFASVQSVSTDSVIN..AQMASQQLPVI
Streptococcus_catagoni 47 SNKYQVNKVSKKKIEKNKEAKGNFDFSSVEAVSTEAVIE..AQMAAQDLPVI
Streptococcus_iniae 24 SNKYQVTKVSKKTIEKNKTAKSSFDFEAVEATSTDSVLE..AQMASQQLPVI
Streptococcus_oralis 47 TNKYQVNQVTKENIDENLKTEGNFDFDSVKSISSEAVLA..SQWDAQKLPVI
Streptococcus_pneumoniae 16 TNKYQVSQVSKEKLEENQDTEGNFDFDSVKAISSEAVLT..SQWDAQKLPVI
Streptococcus_parasanguinis 45 TNQYQVSKVDKNTIDKNKKVKTSFDFQHVKSLSTEAVIN..AQWKAQKLPVI
Streptococcus_anginosus 45 TNRYQVSKVSKDKITQNKKAKTSFNFDKVKSLSTEDVIN..AQWKAQQLPVV
Streptococcus_sp. 45 TNRYQVSNVSKDKINKNKKAKTSFNFDKVKSLSTEDVIN..AQWRAQQLPVI
Streptococcus_suis 45 TNKYQISNVTTEDIEKNKQAETTFDFDQVQSISTEAILA..AQWDAQRLPVI
Streptococcus_minor 45 SNHYQISKVSKEDIEKNKNADVTFDFNQVESISTEAVLK..AQWEAQQLPVI
Streptococcus_azizii 46 SNRYQVTKVTEADIEKNRQAETTFDFEQVESISTEAVLK..AQWEAQELPVI
Streptococcus_cuniculi 53 SNRYQVSKFTEEDLLKNKNAKATFDFEQVNSISTEAVLK..AQWASQELPVI
Streptococcus_macedonicus 95 GGIAVPDVGINLPIFKGLGNTELSYGAGTMKENQVMGGENNYALASHHVFGL
Streptococcus_vicugnae 95 GSIAVPDLGINLPIFKGLGNTELSYGAGTMKEDQVMGGQNNYALASHHVFGV
Streptococcus_equinus 95 GGIAIPDVGINLPIFKGLGNVELSYGAGTMKEEQVMGGQNNYALASHHVFGI
Streptococcus_thermophilus 100 GGIAIPEVGINLPIFKGLGNTELTYGAGTMKENQVMGGENNYSLASHHIFGI
Streptococcus_salivarius 100 GGIAIPEVGINLPIFKGLGNTELTYGAGTMKEDQVMGGENNYSLASHHVFGI
Streptococcus_vestibularis 100 GGIAIPEVGINLPIFKGLGNTELTYGAGTMKENQVMGGKNNYSLASHHVFGI
Streptococcus_ictaluri 96 GGIAIPELGINLPIFKGLGNVELIYGAGTMKEEQVMGGENNYSLASHHIFGM
Streptococcus_phocae 96 GGIAIPELGINLPIFKGLGNVELIYGAGTMKEDQVMGGDNNYSLASHHIFGL
Streptococcus_equi 98 GGIAIPELGINLPIFKGLGNTELIYGAGTMKEDQVMGGENNYSLASHHIFGL
Streptococcus_castoreus 86 GGIAIPDVGINLPIFKGLGNVELMYGAGTMKEDQVMGGDNNYSLASHHIFNM
Streptococcus_pyogenes 96 GGIAIPELGINLPIFKGLGNTELIYGAGTMKEEQVMGGENNYSLASHHIFGI
Streptococcus_dysgalactiae 96 GGIAIPEVGINLPIFKGLGNVELIYGAGTMKEDQVMGGENNYSLASHHIFGI
Streptococcus_canis 96 GGIAIPDVGINLPIFKGLGNVELIYGAGTMKEDQVMGGENNYSLASHHIFGI
Streptococcus_uberis 98 GGIAIPDVGINLPIFKGLGNTELIYGAGTMKENQVMGGDNNYSLASHHIFGM
Streptococcus_bovimastitidis 94 GGIAIPEVSINLPIFKGLGNTELIYGAGTMKEGQVMGGDNNYSLASHHIFGM
Streptococcus_penaeicida 94 GGIAIPEVSINLPIFKGLGNTELIYGAGTMKEGQVMGGDNNYSLASHHIFGM
Streptococcus_didelphis 97 GGIAIPDIGINLPIFKGLGNVELMYGAGTMKENQVMGGDNNYALASHHIFGL
Streptococcus_catagoni 97 GGIAIPEVSINLPIFKGLGNTELIYGAGTMKENQVMGGENNYSLASHHIFGM
Streptococcus_iniae 74 GGIAIPDLDINLPIFKGLGNTELLFGAGTMKEEQVMGGENNYSLASHHIFGM
Streptococcus_oralis 97 GGIAIPEVEINLPIFKGLDNVNLFYGAGTMKPDQKMG.EGNYSLASHHIFTA
Streptococcus_pneumoniae 66 GGIAIPELEMNLPIFKGLDNVNLFYGAGTMKREQVMG.EGNYSLASHHIFGV
Streptococcus_parasanguinis 95 GGISIPELNMNLPIFKGLENVALYYGAGTMKENQVMG.QGNYSLASHHVFGL
Streptococcus_anginosus 95 GGIAIPELSMNLPIFKGLDNVGLYYGAGTMKETQQMG.KGNYALASHHVFGI
Streptococcus_sp. 95 GGIAIPELSMNLPIFKGLDNVGLYYGAGTMKETQQMG.KGNYALASHHVFGI
Streptococcus_suis 95 GGIAVPELGINLPIFKGVFNTSLMYGAGTMKENQEMG.KGNYALASHHIFGV
Streptococcus_minor 95 GGIAIPDLGVNLPIFKGLSNVALMYGAGTMKENQVMG.QGNYTLASHHIFGI
Streptococcus_azizii 96 GGIAIPDLNVNLPIFKGLSNVALMYGAGTMKETQEMG.QGNYSLASHHIFGI
Streptococcus_cuniculi 103 GGIAIPDLNINLPIFKGVSNVSLMYGAGTMKETQEMG.QGNYALASHHIFGI

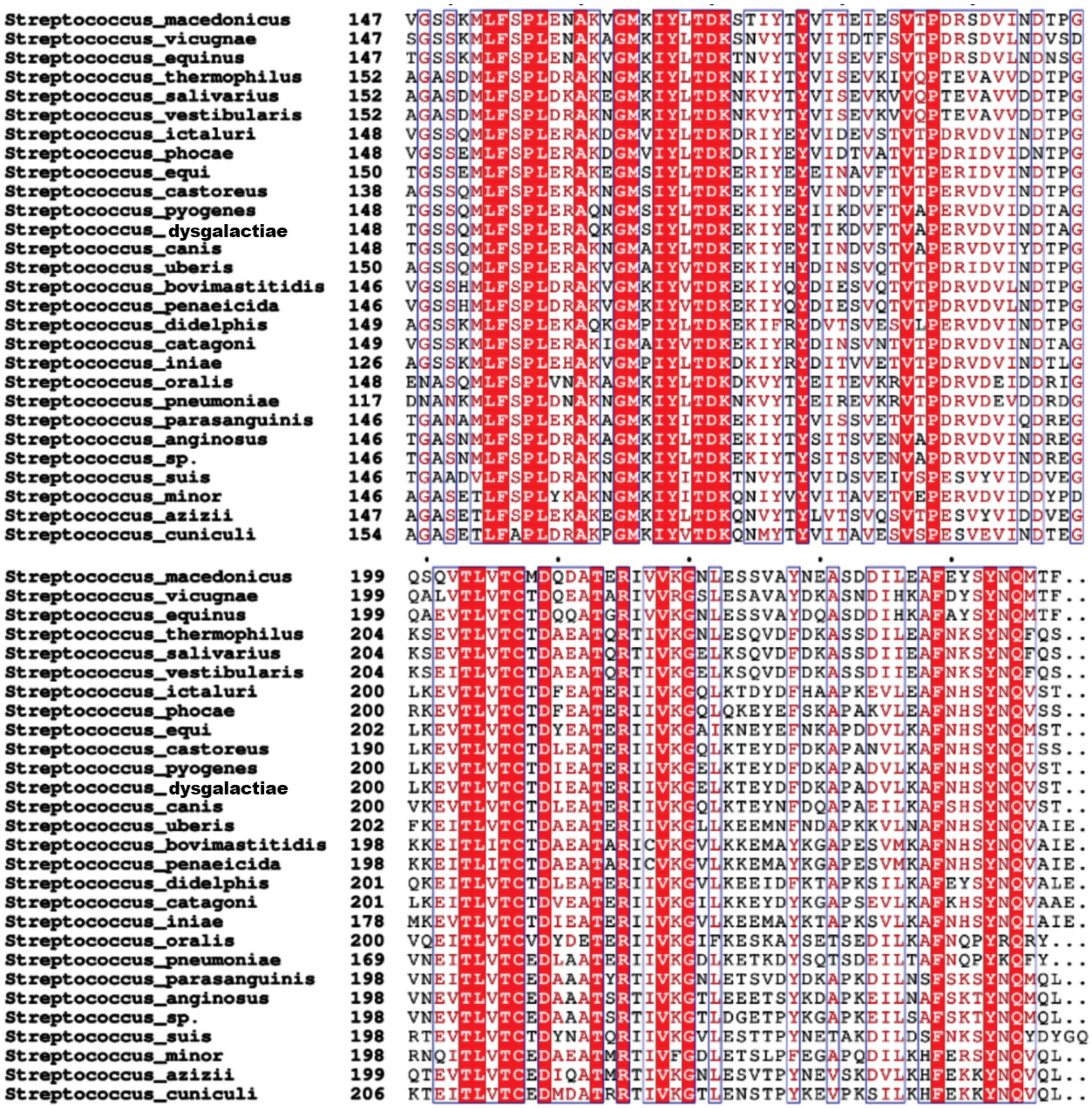


**Figure S6. Multiple sequence alignment of 28 *Streptococcus* SrtA sequences.** Multiple sequence alignment (MSA) was performed using Clustal Omega, as described in the Materials and Methods. The K35 and R38 spySrtA positions are indicated with black arrows.

**Sequences used for AlphaFold2 modeling.**

```
>spySrtA_Q99ZN4_STRP1_1-249
MVKKQKRRKIKSMSWARKLLIAVLLILGLALLFNKPIRNTLIARNSNKYQVTKVSKKQIKKNKEAKSTFDFQAVEPV
STESVLQAQMAAQQLPVIGGIAIPELGINLPIFKGLGNTELIYGAGTMKEEQVMGGENNYSLASHHIFGITGSSQML
FSPLERAQNGMSIYLTDKEKIYEYIIKDVFTVAPERVDVIDDTAGLKEVTLVTCTDIEATERIIVKGELKTEYDFDK
APADVLKAFNHSYNQVST
```

```
>M_protein_M6A_STRP6_376-415
ETKRQLPSTGETANPFFTAAALTVMATAGVAAVVKRKEEN
```
